# Early removal of senescent cells protects retinal ganglion cells loss in experimental ocular hypertension

**DOI:** 10.1101/684811

**Authors:** Lorena Raquel Rocha, Viet Anh Nguyen Huu, Claudia Palomino La Torre, Qianlan Xu, Mary Jabari, Michal Krawczyk, Robert N. Weinreb, Dorota Skowronska-Krawczyk

## Abstract

Experimental ocular hypertension induces senescence of retinal ganglion cells (RGCs) that mimicks events occurring in human glaucoma. Senescence-related chromatin remodeling leads to profound transcriptional changes including the upregulation of a subset of genes that encode multiple proteins collectively referred to as the senescence-associated secretory phenotype (SASP). Emerging evidence suggests that the presence of these proinflammatory and matrix-degrading molecules has deleterious effects in a variety of tissues. In the current study, we demonstrated in a transgenic mouse model that early removal of senescent cells induced upon elevated intraocular pressure (IOP) protects unaffected RGCs from senescence and apoptosis. Visual evoked potential (VEP) analysis demonstrated that remaining RGCs are functional and that the treatment protected visual functions. Finally, removal of endogenous senescent retinal cells after IOP elevation by a treatment with senolytic drug dasatinib prevented loss of retinal functions and cellular structure. Senolytic drugs may have the potential to mitigate the deleterious impact of elevated IOP on RGC survival in glaucoma and other optic neuropathies.

## Introduction

Glaucoma is comprised of progressive optic neuropathies characterized by degeneration of retinal ganglion cells (RGC) and resulting changes in the optic nerve. It is a complex disease where multiple genetic and environmental factors interact^1,2^. Two of the leading risk factors, increased intraocular pressure (IOP) and age, are related to the extent and rate of RGC loss. Although lowering IOP is the only approved and effective treatment for slowing worsening of vision, many treated glaucoma patients continue to experience loss of vision and some eventually become blind. Several findings suggest that age-related physiological tissue changes contribute significantly to neurodegenerative defects that cause result in the loss of vision.

Mammalian aging is a complex process where distinct molecular processes contribute to age-related tissue dysfunction. It is notable that specific molecular processes underlying RGC damage in aging eyes are poorly understood. While no single defect defines aging, several lines of evidence suggest that activation of senescence is a vital contributor^3^.

In a mouse model of glaucoma/ischemic stress, we reported the effects of *p16Ink4a* on RGC death^2^. Upon increased IOP, the expression of *p16Ink4a* was elevated, and this led to enhanced senescence in RGCs and their death. Such changes most likely cause further RGC death and directly cause loss of vision. In addition, the analysis of p16KO mice suggested that lack of *p16Ink4a* gene protected RGCs from cell death caused by elevated IOP^2^. Importantly, elevated expression of *p16INK4a* and senescence were both detected in human glaucomatous eyes^2^. Therefore, for the first time, *p16Ink4a* was implicated as a downstream integrator of diverse signals causing RGC aging and death, both characteristics changes in the pathogenesis of glaucoma. Our findings were further supported by a subsequent report showing that *p16Ink4a* was upregulated by TANK Binding Kinase 1 (TBK1) a key regulator of neuroinflammation, immunity and autophagy activity. TBK also caused RGC death in ischemic retina injury^4^. Of particular note, a recent bioinformatic meta-analysis of a published set of genes associated with primary open-angle glaucoma (POAG) pointed at senescence and inflammation as key factors in RGC degeneration in glaucoma^5^.

Glaucoma remains relatively asymptomatic until it is severe, and the number of affected individuals is much higher than the number diagnosed. Numerous clinical studies have shown that lowering IOP slows the disease progression^6,7^. However, RGC and optic nerve damage is not halted despite lowered IOP, and deterioration of vision progresses in most treated patients. This suggests the possibility that an independent damaging agent or process persists even after the original insult (elevated IOP) has been ameliorated.

We hypothesized that early removal of senescent RGCs that secrete senescent associated secretory proteins (SASP) could protect remaining RGCs from senescence and death induced by IOP elevation. To test this hypothesis, we used an established transgenic p16-3MR mouse model^8^ in which the systemic administration of the small molecule ganciclovir (GCV) selectively kills *p16INK4a*-expressing cells. We show that the early removal of *p16Ink4+* cells has a strong protective effect on RGC survival and visual function. We confirm the efficiency of the method by showing the reduced level of *p16INK4a* expression and lower number of senescent β-galactosidase positive cells after GCV treatment. Finally, we show that treatment of p16-3MR mice with a known senolytic drug (dasatinib) has a similar protective effect on RGCs as compared to GCV treatment in p16-3MR mice.

## MATERIALS AND METHODS

### Animals

All animal experiments were approved by the UC San Diego Institutional Animal Care and Use Committee (IACUC) and adhered to the ARVO Statement for the Use of Animals in Ophthalmic and Vision Research. Adult p16-3MR^8^ or C57BL/6 mice (12-16 weeks old, Jackson Labs) were housed in 20 °C environment with standard (12 h light/dark) cycling, food and water available ad libitum. For all experiments, an equal number of male and female mice were used.

### Drug treatment

The p16-3MR transgenic model (**Figure 1A**), in which the mice carry a trimodal reporter protein (3MR) under the control of p16 regulatory region^8^, allows potent selective removal of senescent cells. The 3MR transgene encodes a fusion protein consisting of Renilla luciferase, a monomeric red fluorescent protein (mRFP) and herpes simplex virus thymidine kinase (HSV-TK) which converts ganciclovir (GCV) into a toxic DNA chain terminator to selectively kill HSV-TK expressing cells. The experimental group of animals was treated by intraperitoneal (IP) administration of GCV (Sigma, 25mg/kg once a day) or dasatinib (Sigma, 5mg/kg) after IOP elevation (see below), and a control group of mice was sham-treated with PBS or vehicle (DMSO). Each mouse underwent unilateral hydrostatic pressure-induced IOP elevation to 90 mm Hg, with the contralateral eye left as an untreated control. The mice were IP injected intraperitonealy with GCV or dasatinib at day 0 (IOP elevation day) and continued for four consecutive days (**Figure 1B**). At day 5, animals were euthanized, retinas were isolated and immunostained with anti-Brn3a antibody to evaluate the number of RGCs. All drugs were prepared according to the UC San Diego Institutional Animal Care and Use Committee (IACUC) standards. To ensure a sterile environment, compounds were prepared under the tissue culture hood using sterile PBS. The final solution was filtered through a 0.22um PES membrane just before injection. Tips, tubes, and syringes were sterile.

**Figure 1.**
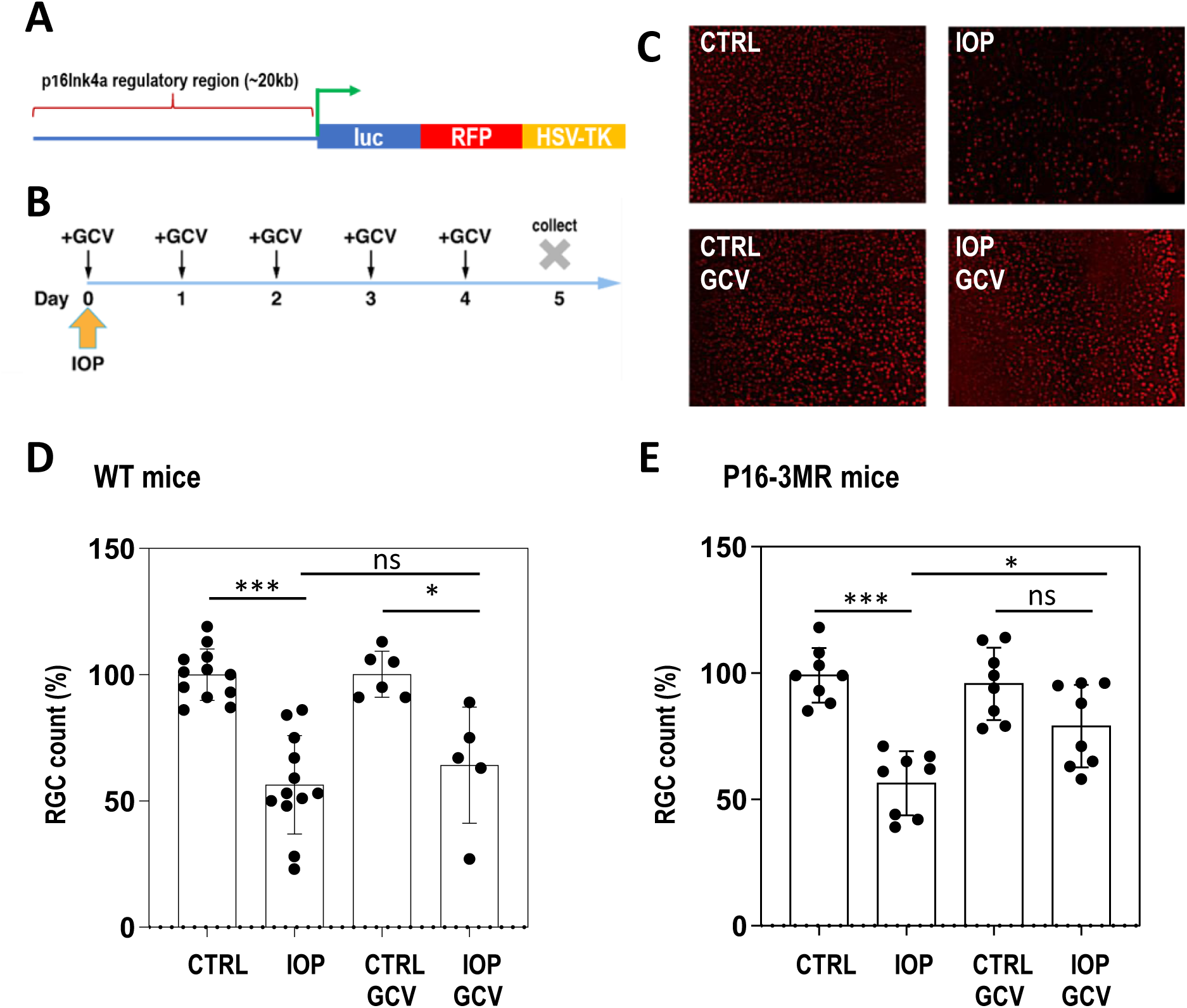
Removal of early senescent cells has a neuroprotective effect on RGCs. **A.** Schematic representation of the p16-3MR transgene. Triple fusion of luciferase, the red fluorescent protein and tyrosine kinase from HSV virus are under control of the regulatory region of p16Ink4a gene. **B.** Plan of the experiment. After unilateral IOP elevation mice are daily injected with GCV (25 mg/kg) intraperitoneally. At day 5 VEP is measured, and tissue is collected for further experiments. **C.** Representative images of retina flat mount immunohistochemistry at day five with anti-Brn3a antibody specifically labeling ∼80% of RGC cells. **D.** Quantification of RGC number at day five after the treatment of WT animals. N>=5 animals in each group **E.** Quantification of RGC number at day five after the treatment of p163MR animals. N=8 animals in each group. In D and E, statistical tests were performed using ANOVA with post-hoc Tukey correction for multiple testing. * p<0.05, ** p<0.01, *** p<0.001, n.s. – not significant

### Hydrostatic intraocular pressure (IOP) elevation

Animals were anesthetized with an intraperitoneal injection of ketamine/xylazine cocktail, (100 mg/kg and 10 mg/kg, respectively), their eyes anesthetized with one drop of proparacaine (0.5 %, Bausch-Lomb) and dilated with one drop of tropicamide (1 %, Alcon Laboratories). Unilateral elevation of IOP was achieved by infusing Balanced Salt Solution (Alcon Laboratories) into the anterior chamber of the eye through using an intravenous (IV) infusion set. The level of IOP increase was determined by the height of the saline bottles on the IV infusion set. Stable elevated IOP of 85-90mm Hg was maintained for 60 minutes, and controlled by IOP measurements using a veterinary rebound tonometer (Tonovet). Both eyes were lubricated throughout testing with an ophthalmic lubricant gel (GenTeal, Alcon Laboratories). Animals recovered on a Deltaphase isothermal pad (Braintree Scientific) until awake. The contralateral eye without IOP elevation served as a healthy non-IOP control (CTRL).

### Visual evoked potential (VEP)

VEP measurements were taken at five days post-IOP elevation. This protocol was adapted from prior studies^9^. Mice were dark adapted for at least 12 hours before the procedure. Animals were anesthetized with ketamine/xylazine and their eyes dilated as above. The top of the mouse’s head was cleaned with an antiseptic solution. A scalpel was used to incise the scalp skin, and a metal electrode was inserted into the primary visual cortex through the skull, 0.8 mm deep from the cranial surface, 2.3 mm lateral to the lambda. A platinum subdermal needle (Grass Telefactor) was inserted through the animal’s mouth as a reference, and through the tail as ground. The measurements commenced when the baseline waveform became stable, 10-15 s after attaching the electrodes. Flashes of light at 2 log cd.s/m^2^ were delivered through a full-field Ganzfeld bowl at 2 Hz. Signal was amplified, digitally processed by the software (Veris Instruments), then exported and peak-to-peak responses were analyzed in Excel (Microsoft). To isolate VEP of the measured eye from the crossed signal originating in the contralateral eye, a black aluminum foil eyepatch was placed over the eye not undergoing measurement. For each eye, peak-to-peak response amplitude of the major component N1 in IOP eyes was compared to that of their contralateral non-IOP controls. Following the readings, the animals were euthanized, and their eyes collected and processed for immunohistological analysis.

### Immunohistochemistry

Following euthanasia, eyes were enucleated and fixed in 4% paraformaldehyde (PFA) in PBS (Affymetrix) for 1 hour and subsequently transferred to PBS. The eyes were then dissected, the retinas flat-mounted on microscope slides, and immunostained using a standard sandwich assay with anti-Brn3a antibodies (Millipore, MAB1595) and secondary AlexaFluor 555 Anti-mouse (Invitrogen, A32727). Mounted samples (Fluoromount, Southern Biotech 0100-01) were imaged in the fluorescent microscope at 20x magnification (Biorevo BZ-X700, Keyence), focusing on the central retina surrounding the optic nerve. Overall damage and retina morphology were also taken into consideration for optimal assessment of the retina integrity. Micrographs were quantified using manufacturer software for Brn3a-positive cells in 6 independent 350 µm × 350 µm areas per flat-mount.

### Real-Time PCR

Total RNA extraction from mouse tissues, cDNA synthesis, and RT-qPCR experiments were performed as previously described^2^. Assays were performed in triplicate. Relative mRNA levels were calculated by normalizing results using GAPDH. The primers used for RT-qPCR are the same as in^2^. The differences in quantitative PCR data were analyzed with an independent two-sample t-test.

### SA-βgal Assay to Test Senescence on Retinas Mouse Eyes

Senescence assays were performed using the Senescence b-Galactosidase Staining Kit (Cell Signaling) according to the manufacturer’s protocol. Images were acquired using a Hamamatsu Nanozoomer 2.0HT Slide Scanner and quantified in independent images of 0.1 mm^2^ covering the areas of interest using Keyence software.

### RNA-Seq analysis

High-quality RNA was extracted using TRIzol Reagent (Invitrogen) and treated with TURBO DNA-free Kit (Invitrogen). RNA-Seq libraries were made from 1 µg total RNA per tissue sample using TruSeq stranded mRNA Library Prep Kit (Illumina, kit cat. no. 20020597) according to the manufacturer’s instructions. The libraries were size selected using Agencourt Ampure XP beads (Beckman Coulter) and quality checked by Bioanalyzer (Agilent). The strand-specific RNA-Seq paired-end reads sequence data (PE50: 2×50 bp) were obtained on a HiSeq4000. RNA-Seq reads were counted using HOMER software considering only exonic regions for RefSeq genes.

## RESULTS

Intraocular pressure was increased in one eye of transgenic mice bearing the p16-3MR construct (**Figure 1A**). After IOP elevation, mice were intraperitonealy injected with GCV for five consecutive days (**Figure 1B)** to specifically deplete p16Ink4a-positive (p16^+^) cells. In parallel, wild-type animals were subjected to the same protocol, i.e., underwent five daily GCV injections after unilateral IOP elevation. Retina flat mount immunohistochemistry and RGC quantification were used to assess potential impact of drug treatment. We observed that five-day administration of GCV after IOP elevation had a significant protective effect on Brn3a^+^ RGC number in p16-3MR mice when compared to untreated eye (**Figure 1C, D**). This protection was not observed in WT animals (**Figure 1E**) confirming that the effect of the GCV injection is specific to the mice harboring the GCV-sensitive transgene.

Next, to test whether the protection of RGC numbers in GCV treated retinas was accompanied by the protection of the the visual circuit integrity on day five, the *in vivo* signal transmission from the retina to the primary visual cortex was assessed by measuring visual evoked potentials (VEP) (**Figure 2A**)^10,11^. In brief, the reading electrode was placed in the striate visual cortex, with the reference electrode in the animal’s mouth and ground electrode in the tail. Flash stimuli were presented in a Ganzfeld bowl. Response amplitudes were quantified from the peak-to-peak analysis of the first negative component N1. Using this approach, we have found that eyes subjected to IOP elevation showed decreased VEP P1-N1 amplitude (**Figure 2B**), compared to contralateral non-IOP control eyes. However, there was a marked rescue of VEP signals in transgenic animals treated with GCV (**Figure 2B).** Further quantification showed significant vision rescue upon GCV treatment only in p16-3MR and not WT animals (**Figure 2C, D**); confirming the specificity of GCV treatment.

**Figure 2.**
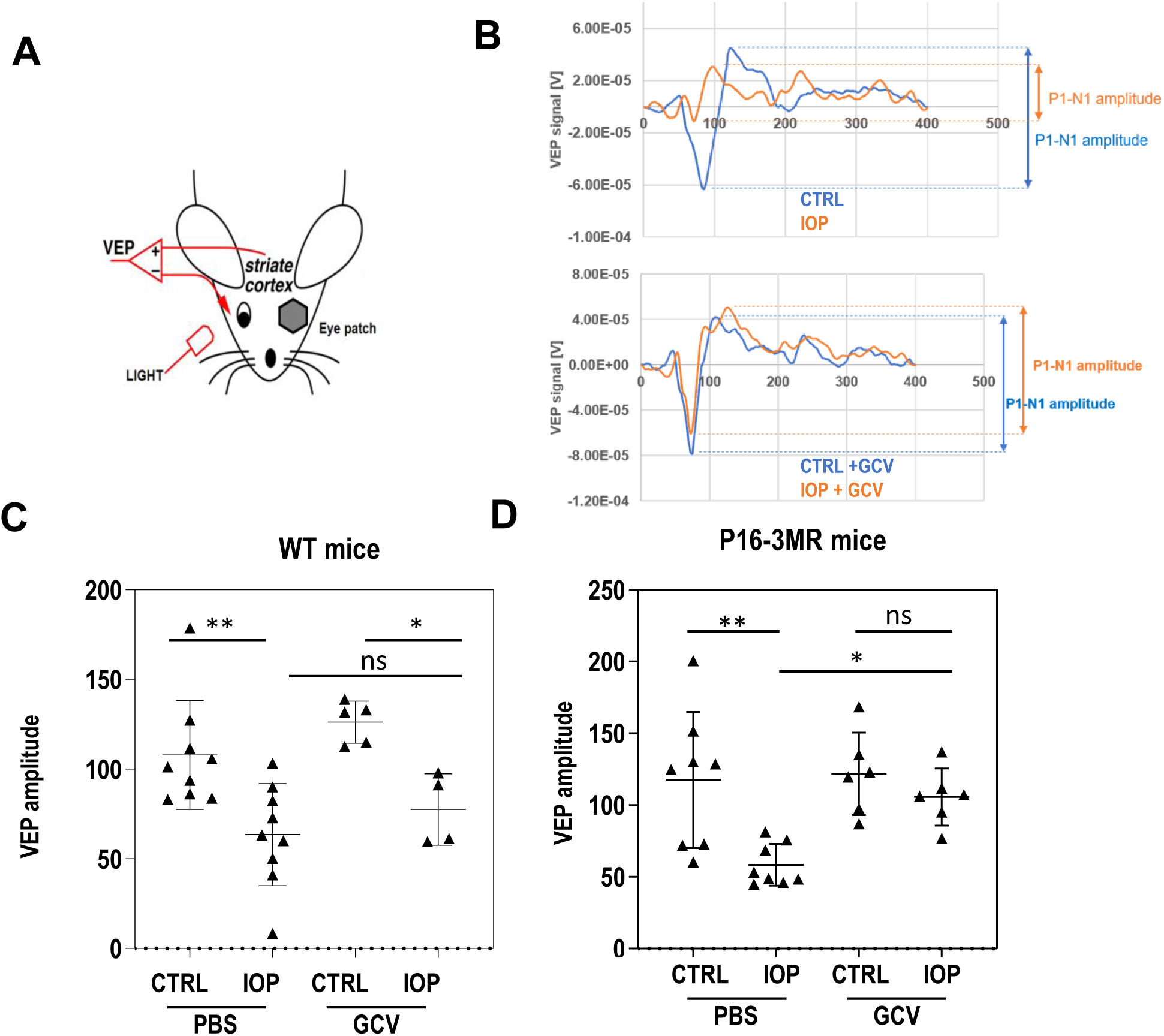
Visual functions are preserved in animals when senescent cells were removed. **A.** Schematic representation of the placement of electrodes for VEP measurements. **B.** Example results of VEP experiments. Top: results of the VEP response of healthy and IOP treated eyes. Bottom: After GCV injections the VEP response is protected. **C.** Quantification of VEP responses at day five after the treatment of WT animals. N>=4, **D.** Quantification of VEP responses at day 5 after the treatment of the p16-3MR animals. N>=6. In C and D, statistical tests were performed using ANOVA with post-hoc Tukey correction for multiple testing. * p<0.05, ** p<0.01, *** p<0.001, n.s. – not significant

Our previous studies indicated that the increase in *p16INK4a* expression could be first observed as early as day three post-IOP elevation^2^. Therefore, we chose this time-point to analyze the effectiveness of GCV treatment on senescent cells in treated and control retinas of p16-3MR animals. RGC quantification showed that in animals not injected with GCV only ∼15-20% of cells disappeared at day 3 (compared to ∼45-50% on day 5).

To test whether GCV treatment indeed removed senescent cells in the retina we used two approaches. First, we quantified the *p16Ink4a* expression at day three post IOP treatment in GCV treated and control retinas. Expectedly, GCV treatment prevented IOP-induced increase in *p16Ink4a* expression observed in non-treated eyes (**Figure 3B**). Importantly, this was accompanied by significant decrease in numbers of IOP-induced β-galactosidase-positive senescent cells in 3-day GCV-treated retinas as compared to non-treated cells (**Figure 3C, D**). This indicates that IOP-induced early senescent cells are efficiently removed by GCV-treatment by day 3, what precedes the RGC loss observed in non-treated eyes between day 3 and day 5.

**Figure 3.**
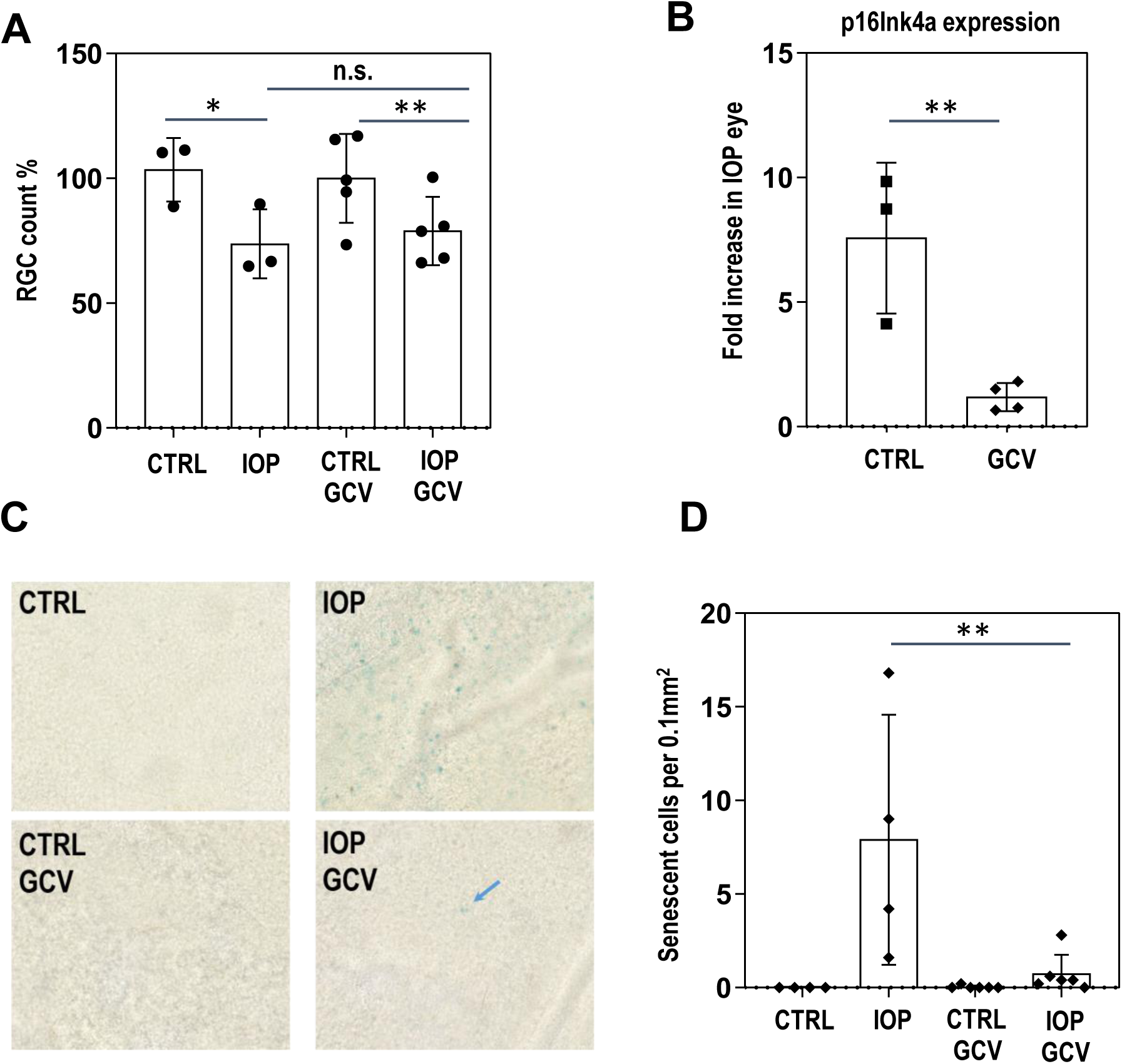
Senescence is lowered upon GCV treatment ∼2 days before the effects on RGC numbers are observed. **A.** At day 3 after IOP only 20% of RGCs are lost compared to the non-treated eye. Similar numbers of cells are lost in GCV treated eyes at this stage. N=3 (non-GCV) and N=5 (GCV), ANOVA, *p<0.05, **p<0.01, n.s. – not significant **B**. p16Ink4a expression is significantly lower in affected retinas isolated from GCV treated p16-3MR animals at day 3 after IOP elevation. T-test, **p<0.01 **C**. Number of SA-b-gal positive cells is lowered upon GCV treatment. Blue arrow – remaining senescent cell **D**. Quantification of number of senescent cells upon IOP elevation in retinas isolated from mouse treated and non-treated with GCV. N=4 (non-GCV), N=6 (GCV); ANOVA, **p<0.01

We next set forward on trying to understand molecular changes underlying the apparent protective effect of the removal of senescent cells by GCV treatment in p16-3MR animals. First, we performed time-course experiment in wild-type mice to follow the activation of caspase expression as a marker of endogenous stress in the cell (**Figure 4A**). We quantified both total number of RGC (by nuclear staining for RGC specific transcription factor Brn3a) and activated caspase-positive cells (by phospho-caspase3 staining). The highest number of RGCs with concomitant staining of activated caspase was observed at day 3 after IOP elevation (**Figure 4A** *right*). As shown above, at day 3 most of the RGCs are still present (**Figure 3A**). Day 3 also corresponds to the highest expression levels of senescence associated factors, as previously observed^2^. We thus reasoned that relevant effects of GCV treatment should be easy to observe at around this stage. To identify potential differences in an unbiased way, we performed RNA-seq analysis in IOP and non-IOP retinas with or without GCV treatment. Total RNA isolated from 3 retinas in each experimental group was converted to cDNA libraries and sequenced. Of the total 21351 detected gene loci, 1601 were significantly de-regulated by the IOP treatment; 999 detected gene loci were up-regulated and 602 down-regulated (**Figure 4B, *top***). When the IOP treatment was performed in mice treated with GCV, the total numbers of IOP-affected genes changed modestly to 1707, with 848 up-regulated and 859 down-regulated (**Figure 4B, *bottom***).

**Figure 4.**
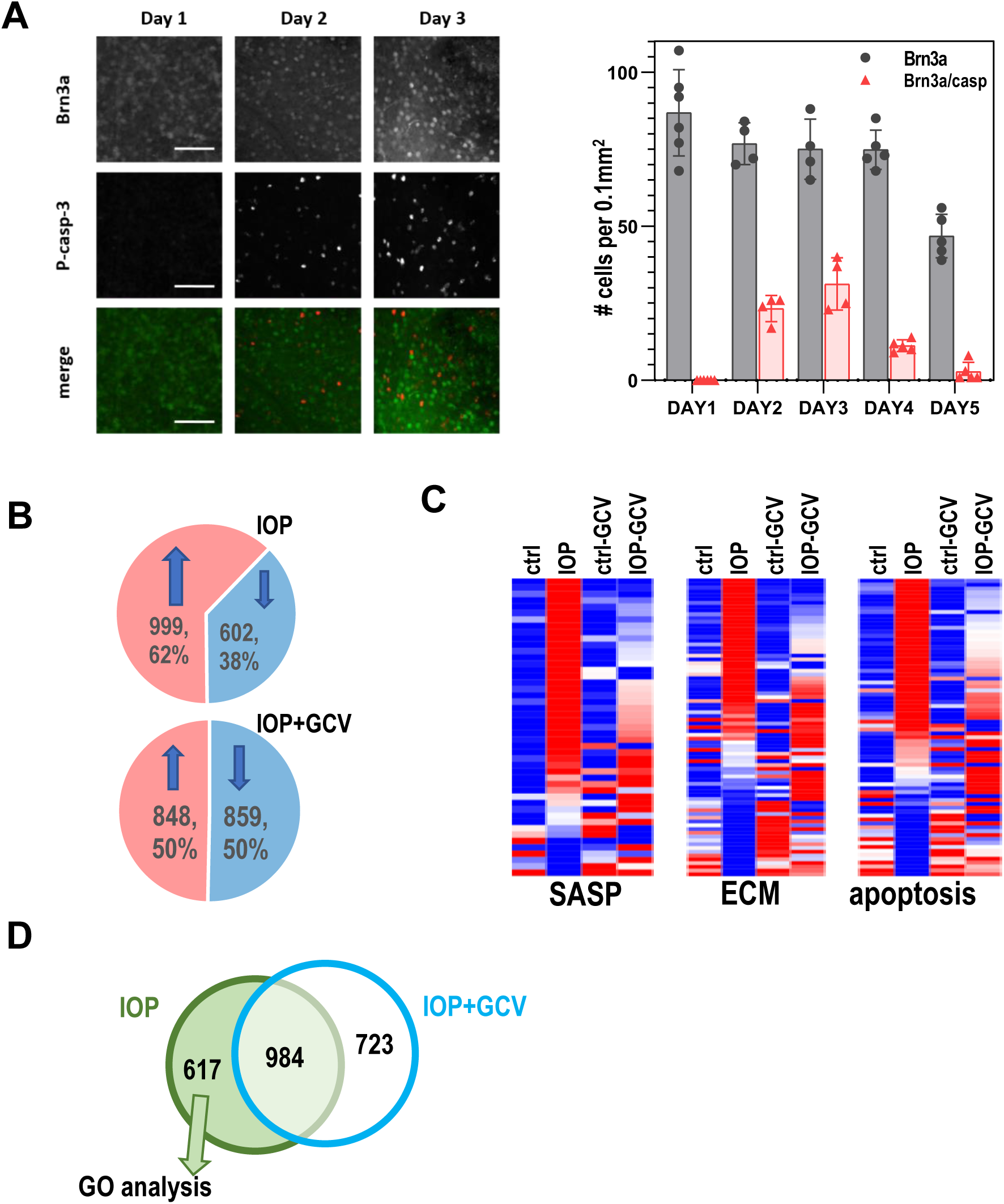
Analysis of pathways affected in IOP treated retinas. **A.** Immunohistochemistry of Brn3a and activated caspase show increase of apoptosis at day1, 2 and 3 after IOP treatment. left: quantification of the time-course experiments followed by immunochemistry with Brn3a and activated caspase 3; **B**, RNA-seq analysis of response to IOP and GCV. Eyes subjected to IOP elevation show significant change in gene expression with more genes upregulated than downregulated. Similar analysis in GCV treated animals shows close to equal distribution of upregulated and downregulated genes **C.** Heatmap analysis of genes involved in senescence, active oxidative species (ROS), apoptosis, extracellular matrix homeostasis (ECM) and inflammation. **D.** Venn diagram showing overlap between genes dysregulated upon IOP in GCV treated and untreated retinas. 617 genes specifically dysregulated in IOP only retinas were used for GO analysis.

To inquire in an unbiased way about the differences in signaling pathways and cellular processes affected by IOP, GO analysis using PANTHER was performed^12^. This approach revealed that processes of the immune system response, inflammation, and extracellular matrix composition and cell-matrix interaction were significantly changed in IOP samples (**Table 1**). We have also detected the significantly deregulated genes involved in apoptosis, microglial activation and interlukin-6 and 8 production and secretion. This analysis shows that many mechanisms are induced upon an acute IOP elevation, most probably causing additional transcriptional stress to cell.

Further analysis revealed that the genes involved in cellular senescence, extracellular matrix molecules and in factors involved in apoptosis (**Table 2**)^13^ were significantly de-regulated upon IOP elevation. Importantly, 3-day treatment to remove p16+ cells significantly mitigated this response (**Figure 4C**). This data is in agreement with the loss of the senescence cells upon GCV treatment (**Figure 3B, C, D**) and lower detrimental impact of senescent cells on surrounding cells.

Additional GO analysis of the 617 genes which were significantly de-regulated upon IOP elevation specifically in non-treated retinas (ie. genes where the effects of IOP were dampened by GCV-mediated removal of senescent cells) (**Figure 4D)** identified a specific enrichment of a class of genes belonging to the ABL1 pathway and ABL1 downstream targets (**Fig S1**). Prompted by this finding, we explored whether dasatinib, a well known senolytic drug and a Bcr-Abl and Src family threonine kinase inhibitor, could have a beneficial effect similar to GCV in p16-3MR mice. To this end, p16-3MR mice were treated with dasatinib (5mg/kg) or vehicle for 5 days by intraperitoneal injection, similarly to the experimental procedure used for GCV (**Figure 1B**). Performing this experiment in the transgenic mice allowed direct comparison of the efficiencies of both treatments in the same mouse strain. At day five after IOP elevation, VEP measurement was performed and retinas were immunostained to quantify RGC loss. We observed that dasatinib treatment prevented the loss of RGC (**Figure 5C**) similar to what was observed in GCV treated animals (**Figure 1E**). Most importantly, VEP analysis revealed that senolytic drug treatment successfully prevented vision loss upon IOP elevation (**Figure 5D**).

**Figure 5.**
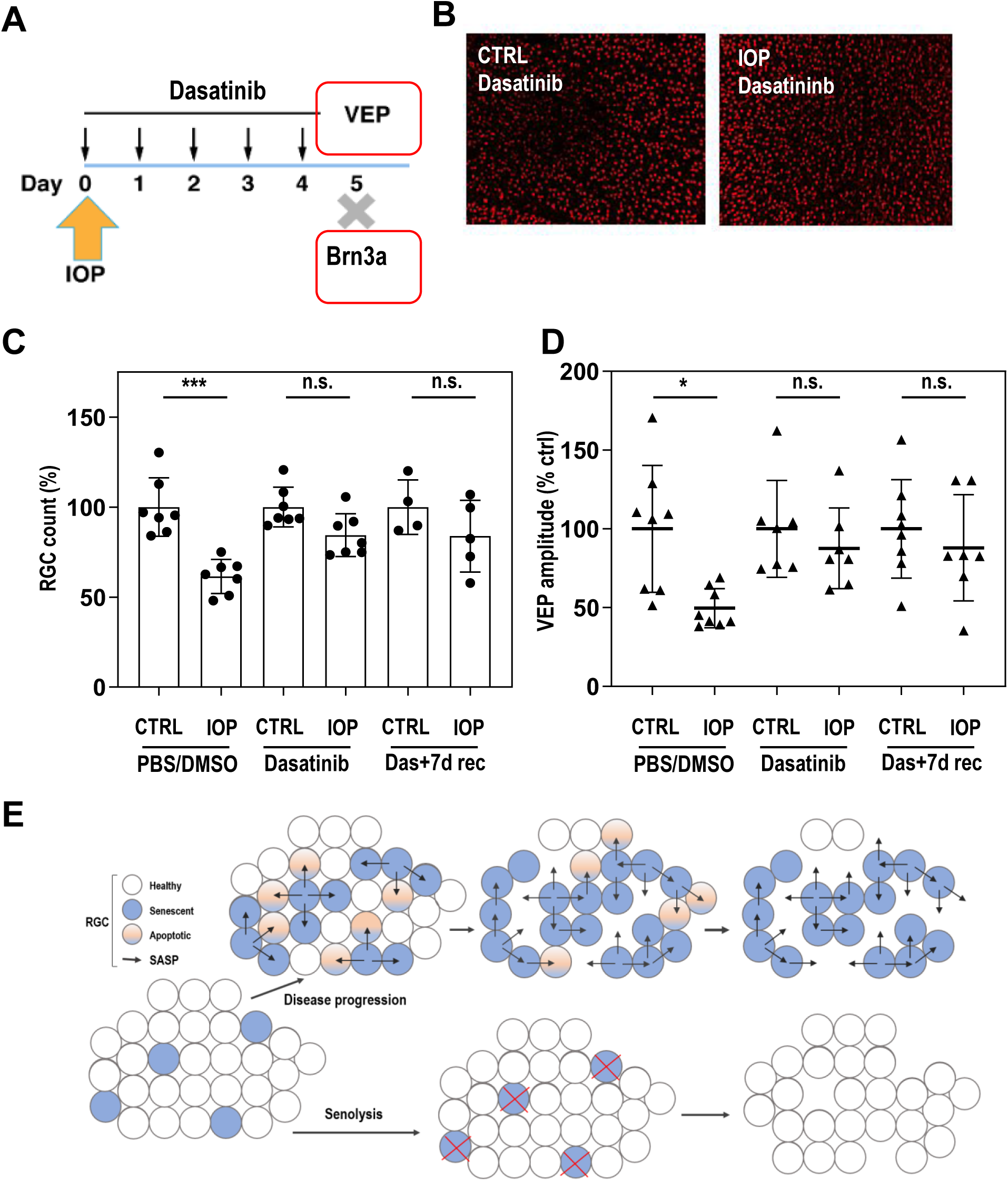
Dasatinib protects retina degeneration. **A.** plan of the experiment. After unilateral IOP elevation mice are daily injected with dasatinib (5 mg/kg) intraperitoneally. At day 5 VEP is measured and tissue is collected for further experiments. Immunohistochemistry of Brn3a and activated caspase show increase of apoptosis at day 3 after IOP treatment. **B**, Retina flat mount immunohistochemistry at day 5 with anti-Brin3a antibody specifically labelling ∼80% of RGC cells. **C.D.** Quantification of RGC number (C) or VEP responses (D) at day 5 (four conditions) or day 12 (additional 7 days of “recovery”, two conditions) after the 5d ay treatment of p16-3MR animals with dasatinib. N>4 animals in each group. Statistical tests were performed using ANOVA with post-hoc Tukey correction for multiple testing. * p<0.05, *** p<0.001, n.s. – not significant **E.** Model. <u>Top</u>: Upon elevated IOP damaged cells become senescent and start to express SASP molecules. While disease progresses the SASP molecule induce senescence or apoptosis in neighboring cells. <u>Bottom</u>: When senescent cells are removed using senolytic drug the neighboring cells are not exposed to detrimental SASPs and the disease progression is significantly slowed down. Remaining cells are healthy.

Finally, we explored whether the protective impact of the drug is caused by the sustained inhibition of the cellular processes and whether it is maintained even after the drug is no longer present. To do that, p16-3MR mice were treated with dasatinib (5mg/kg) or vehicle for 5 days by intraperitoneal injection, similarly to the experimental procedure used for GCV (**Figure 1B**). After that, the mice were no longer treated with drug or PBS and at day twelve after IOP elevation, functional measurement was performed and RGCs were quantified. Also in this treatment regime, dasatinib prevented the loss of RGC (**Figure 5C**) similar to what was observed in GCV treated animals (**Figure 1E**). Additionally, VEP analysis revealed that senolytic drug treatment with seven days “chase” still successfully prevented vision loss upon IOP elevation (**Figure 5D**).

## DISCUSSION

The collective findings of the current study strongly support the notion that removal of senescent cells provides beneficial protective effect to retinas damaged by elevated IOP. Here, we show that in transgenic animals expressing viral TK under the control of regulatory regions of p16Ink4a^8^, selective removal of early senescent cells in the retina is beneficial for neighboring cells undergoing cellular stress induced by IOP elevation. In this model, the treatment with GCV selectively induces cell death of transgene-expressing cells. Early application of GCV as soon as day 0 after the IOP elevation and followed for five consecutive days ensures early removal of *p16Ink4a* expressing cells resulting in protection of neighboring RGCs from death. Remaining cells are still able to provide a signal to the visual cortex, as evidenced by VEP measurements, demonstrating that protected cells are functional.

Three days after GCV injection there is a significant drop in senescent cell number and an accompanying alteration of the transcriptional programs in remaining cells as compared to retinas from untreated mice. Using an RNA-seq approach, we noted significant changes in the senescence program as well as in the extracellular matrix (ECM) function. Both pathways are downregulated by GCV treatment. Finally, we show that dasatinib, a known senolytic drug, can be used to protect RGCs from death, further confirming that early removal of senescent cells induced upon IOP elevation protects retina health. The observation also was confirmed when the RGC count and VEP were assessed seven days after the treatment was stopped; this suggests that the impact of the drug is not reversible during this time. Further studies should investigate different regimes and dosages of the senolytic drugs and their neuroprotective role.

Taken together, the results prompt us to propose a model of how increased IOP leads to the destrucion of retinal structures during glaucoma progression (**Figure 5E**). During early stages, elevated IOP induces cell-intrinsic changes leading to cellular senescence and production of SASP. As the disease worsens, SASP molecules induce apoptosis and senescence in neighboring cells. Such changes are largely independent of whether the initial insult is still present or has been eliminated with IOP-lowering treatment. RGC apoptosis inevitably leads to the loss of axons and optic nerve degeneration. Conversely, when senescent cells (induced directly by elevated IOP) are removed using senolytic drugs (**Figure 5E, *bottom***), neighboring cells are not exposed to detrimental SASP and remain healthy. We propose that such treatment can lead reduce the rate of glaucoma worsening. Moreover, we speculate that changes in combination with IOP-lowering treatments may have even better protection than either type of therapy alone.

We and others previously used 90 mm Hg as an extremely acute and reproducible way to induce cell response and RGC death^2^. This level of pressure is likely an ischemic insult. The resulting synchronized and quick cell death provide unique qualities that are extremely useful for molecular and biochemical studies, however such an acute and high pressure change is not fully representative of POAG, a chronic optic neuropathy. However, this acute insult allows the study of stress response time course and can help unravel important aspects of stepwise response of retinal cells to elevated IOP which is a daunting task to assess in chronic models the disease. Importantly, our previous data showing the presence of senescent cells in human glaucomatous retinas^2^ should stimulate the use of senolytic drugs in other animal models of glaucoma.

Dasatinib is a selective tyrosine kinase receptor inhibitor that is commonly used in the therapy of chronic myelogenous leukemia (CML). Other studies have shown that treatment with dasatinib is effective in destroying senescent fat cell precursors^14^. Our RNA-seq data pointed to this senolytic drug as a potential candidate for in-vivo treatment of retinal damage induced by IOP elevation. Notably, we found that the level of RGC protection resembles the one obtained with GCV treatment of p16-3MR transgenic line. Based on these findings, we conclude that dasatinib treatment resulted in RGC protection through removal of senescent cells. It will be of interest to further investigate the possible therapeutic effects of other senolytic drugs in glaucoma and glaucoma models.

The gene encoding *p16INK4a, CDKN2A*, lies within the INK4/ARF tumor suppressor locus on human chromosome 9p21; this is the most significant region to be identified as having an association with POAG in different population samples^15^. Although the molecular mechanism of many of these associations is yet to be described, we have shown that one of them especially highly correlates with the presence of another top risk variant of glaucoma – Six6 rs33912345. Our study further showed that upregulation of homozygous *SIX6* risk alleles (CC) leads to an increase in *p16Inka* expression, with subsequent cellular senescence^2^. Interestingly, others have described an alternative mechanism whereby IOP induced TBK1 expression caused an increase of *p16Ink4a* expression through the Akt-Bmi1 phosphorylation pathway^4^. Given the complexity of the 9p21 locus, we believe that there are more pathways involved in *p16Ink4a* regulation and further work is needed to understand the role of p16Ink4a as a integrator of these signals especially upon IOP elevation.

Several collaborative efforts identified numerous SNPs localized within the 9p21 locus to be highly associated with the risk of open-angle glaucoma including Normal-Tension Glaucoma (NTG), a glaucomatous optic neuropathy not associated with elevated IOP^16,17^. Intriguingly, one of the top variants associated with the risk of NTG is located in the gene TBK1, a factor that has been recently shown to be implicated in upregulation of *p16ink4a* gene^4^. Finally, recent studies have also revealed that specific methylation patterns in the 9p21 locus are strongly associated with the risk of NTG glaucoma^18^. It is notable that the positions of most, if not all, of these SNPs and methylation markers, overlap with active regulatory regions within the locus identified by ENCODE^19^. Although regulation of the 9p21 locus in the context of many diseases and aging is under extensive investigation, it still remains to be explicitly addressed in relation to glaucoma.

Another major type of glaucomatous optic neuropathy is angle closure glaucoma (ACG), a condition characterized by blockage of the drainage angle of the eye. To date, there is no study reporting genetic variants or methylation markers in the 9p21 locus significantly associated with the risk of ACG despite several studies implicating various molecular mechanisms ^20^. Nevertheless, the fact that progressive vision loss is observed in PACG patients, even after lowering the IOP^21^, raises the question whether an association could be observed between 9p21 markers and the *progression* rather than the *risk* of the disease. Further studies to unravel such associations are necessary.

Markers of cellular senescence such as expression of the *p16Ink4a* and SASP molecules dramatically increase during aging in both humans and mice. Several studies suggest that *p16Ink4a*+ cells act to shorten healthy lifespan by promoting age-dependent changes that functionally impair tissues and organs^22-25^. Intriguingly, a recent explosion of studies has shown that removal of senescent cells using senolytic drugs in progeroid (accelerated aging phenotype) and healthy mice induces lifespan extension and improves the health of animals^22,26-28^. Our studies suggest a potential use of such therapy to reduce glaucoma associated blindness, either as a stand-alone treatment or together with IOP-lowering therapies.

## DATA AVAILABILITY STATEMENT

The data that support the findings of this study are openly available in GEO database (GEO number pending) and in Dryad at https://doi.org/10.6075/J0707ZTM

## ACKNOWLEDGEMENTS

We thank Sherrina Patel for help with this project. This work was supported by R01 EY02701, RPB Special Scholar Award and Glaucoma Research Foundation Shaffer Award to D.S.K. as well as by RPB Unrestricted Grant to Shiley Eye Institute. Functional imaging and histology work were funded by the UCSD Vision Research Center Core Grant P30EY022589.

## FIGURE LEGENDS

**Supplementary figure 1.**
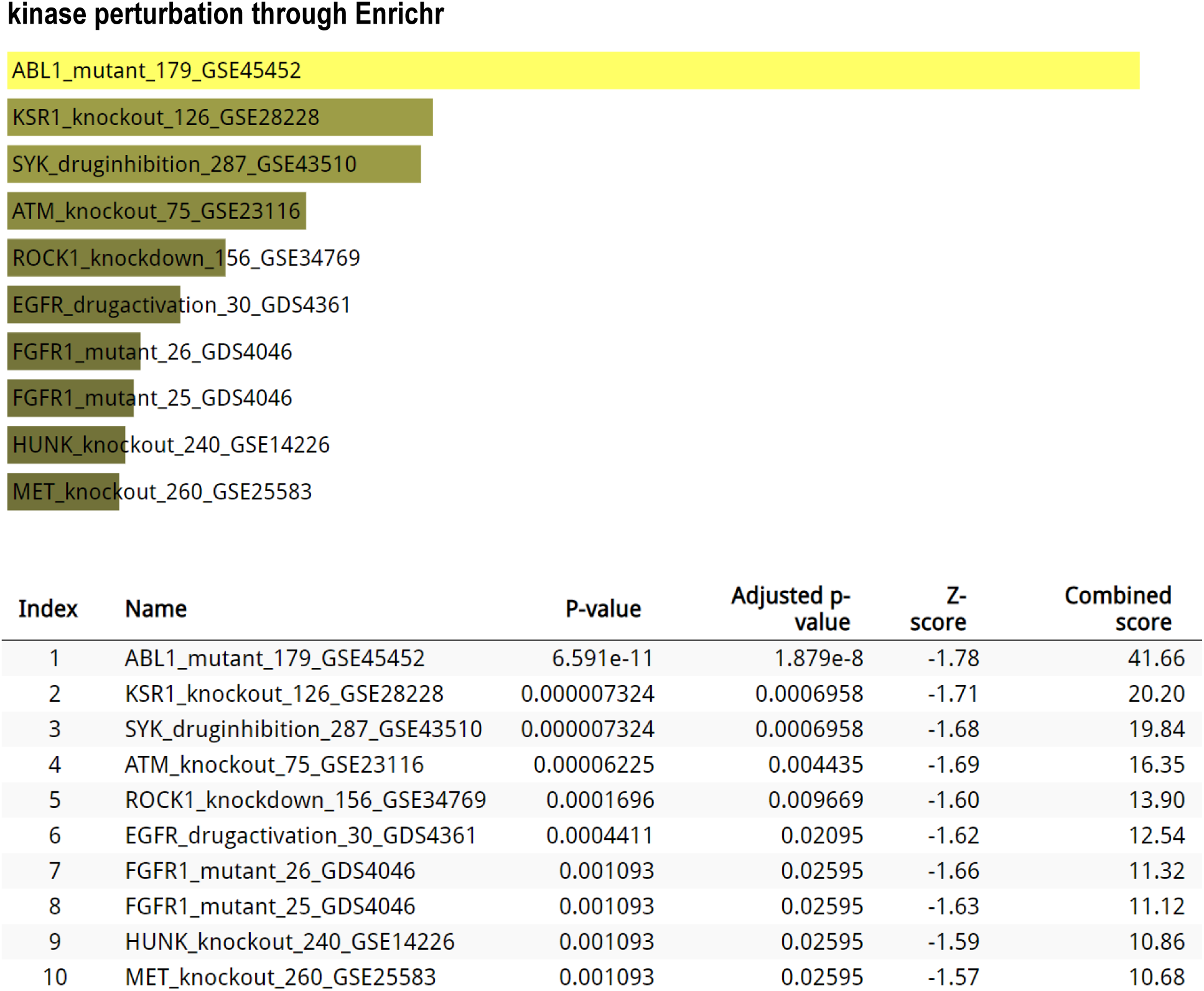
Enrichr analysis of genes. 617 genes altered in nGCV conditions by IOP.

**Supplementary figure 2.**
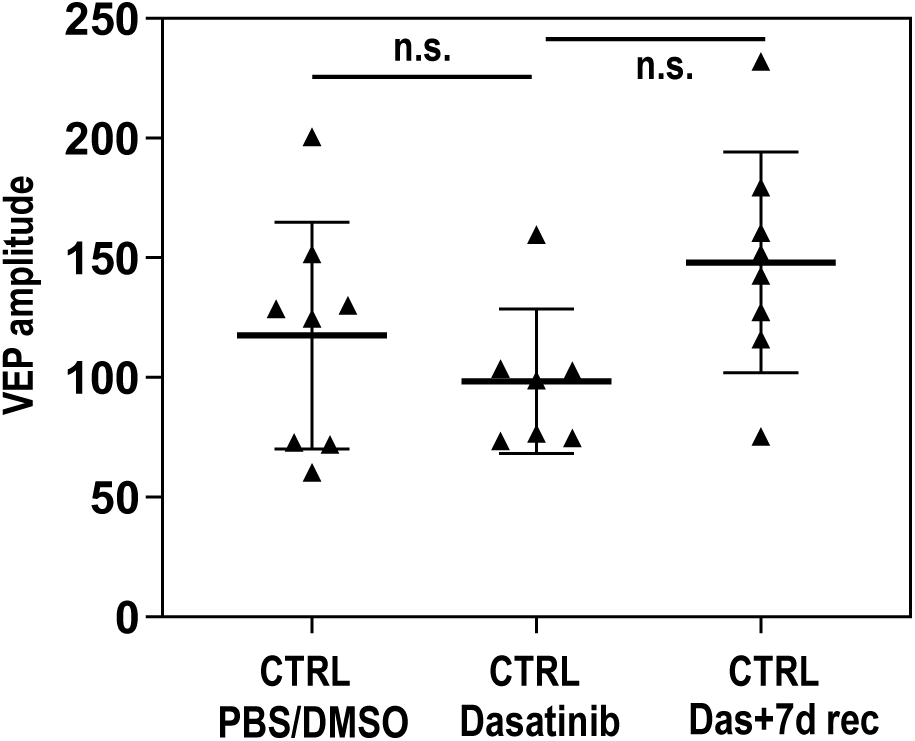
Comparison of the healthy eye VEP amplitude. No significant differences detected. N>4 animals in each group. Statistical tests were performed using ANOVA with post-hoc Tukey correction for multiple testing. n.s. – not significant

---

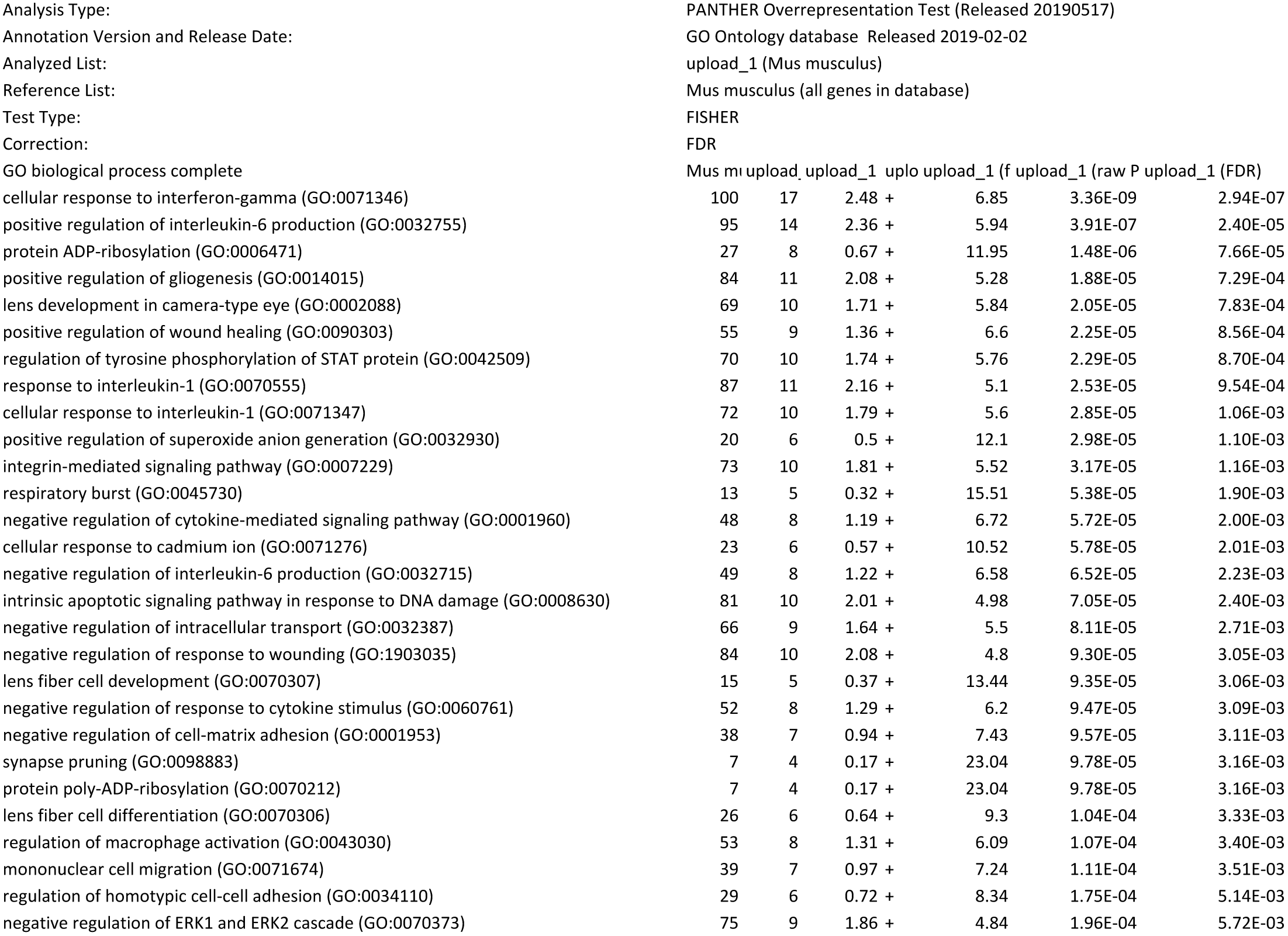
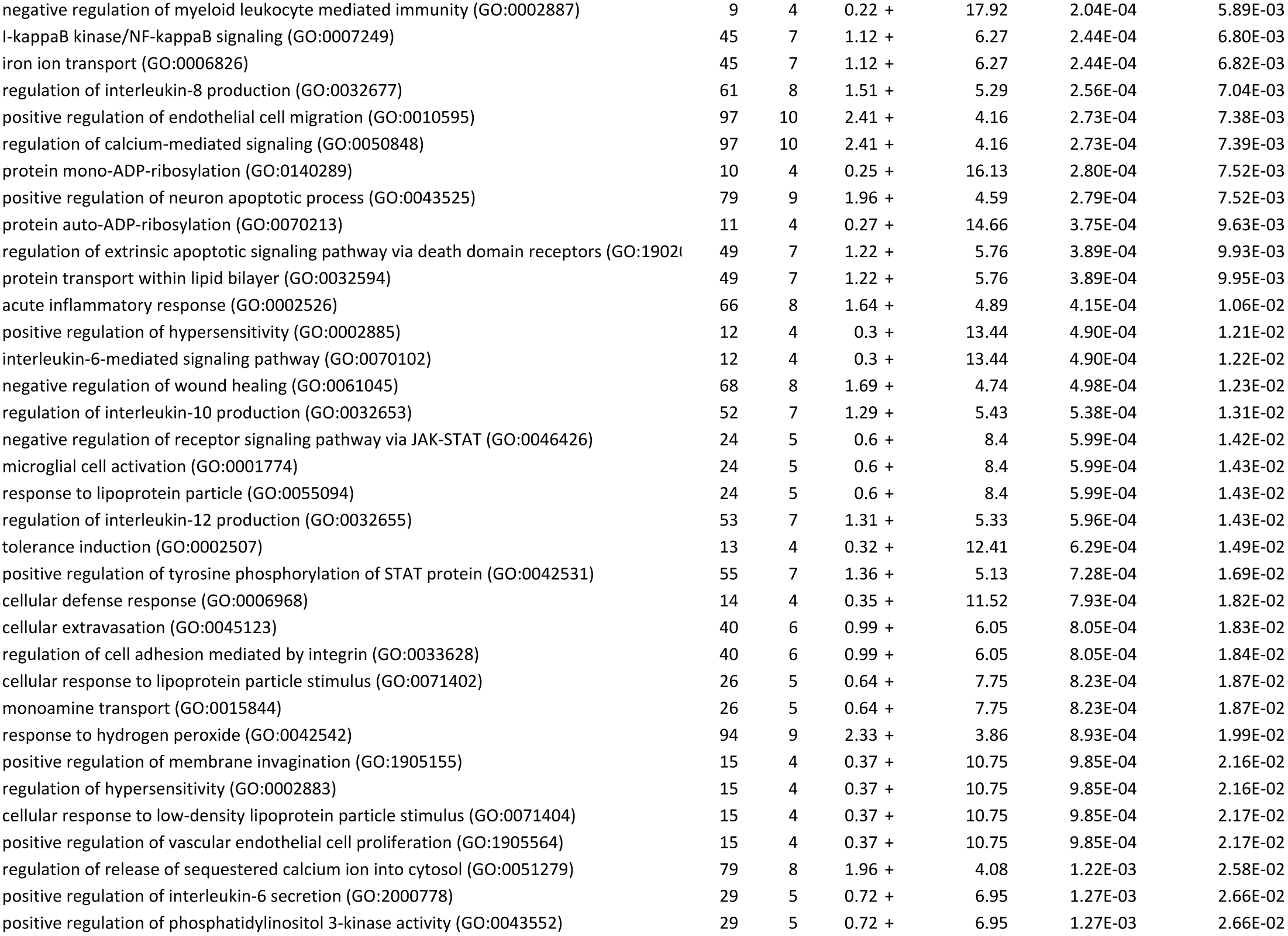
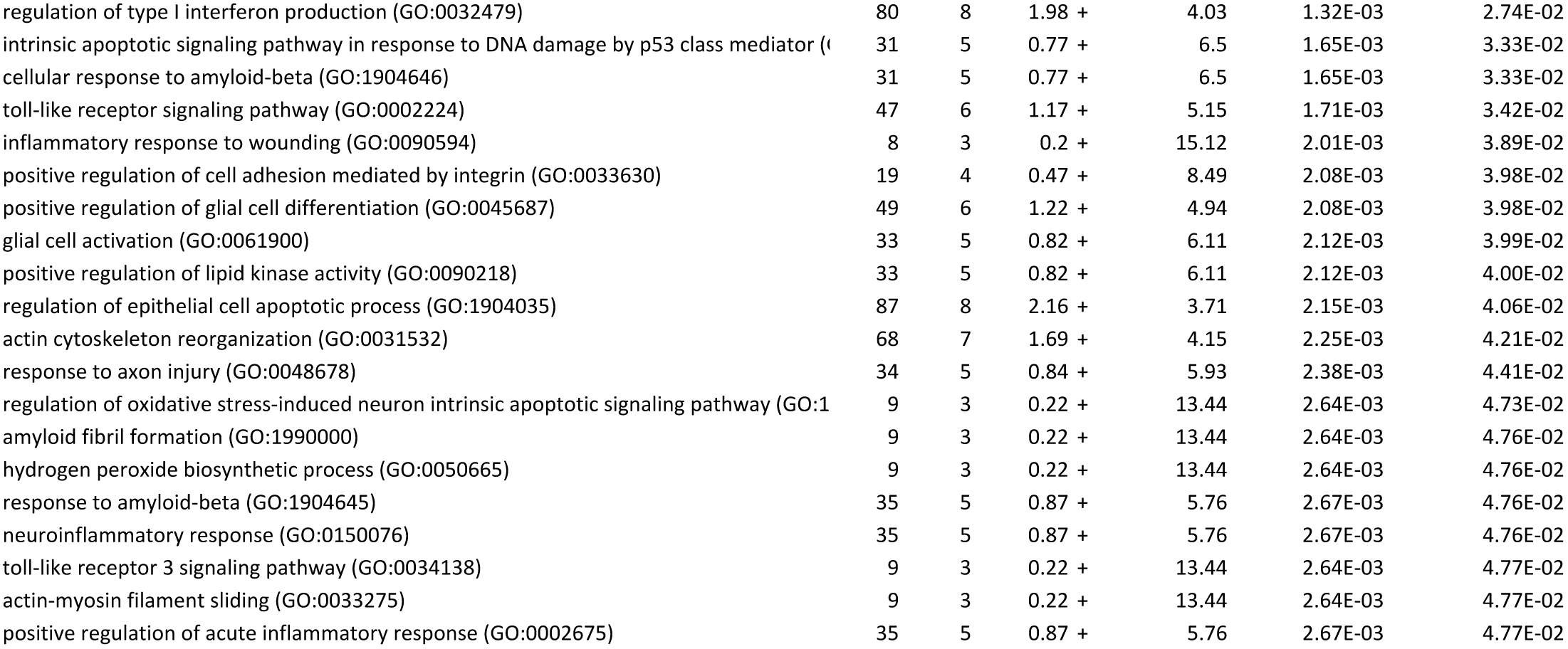

| Apoptosis |  |
| --- | --- |
| Abl1 | Gadd45a |
| Actb | Gapdh |
| Aifm1 | Gusb |
| Akt1 | Hsp90ab1 |
| Anxa5 | Igf1r |
| Apaf1 | Il10 |
| Api5 | Lhx4 |
| Atf5 | Ltbr |
| Bag1 | Mapk1 |
| Bag3 | Mcl1 |
| Bak1 | Naip1 |
| Bax | Naip2 |
| Bcl10 | Nfkb1 |
| Bcl2a1a | Nme5 |
| Bcl2l10 | Nod1 |
| Bcl2l2 | Nol3 |
| Bid | Polb |
| Birc2 | Prdx2 |
| Birc3 | Ripk1 |
| Birc5 | Tnfrsf10b |
| Bnip2 | Tnfrsf11b |
| Bok | Tnfrsf1a |
| Card10 | Tnfsf10 |
| Casp1 | Tnfsf12 |
| Casp12 | Traf1 |
| Casp14 | Traf2 |
| Casp2 | Trp53 |
| Casp3 | Trp53bp2 |
| Casp4 | Trp63 |
| Casp6 | Trp73 |
| Casp7 | Xiap |
| Casp8 |  |
| Casp9 |  |
| Cd40lg |  |
| Cd70 |  |
| Cflar |  |
| Cidea |  |
| Cideb |  |
| Cradd |  |
| Dad1 |  |
| Dapk1 |  |
| Dffa |  |
| Dffb |  |
| Diablo |  |
| Fas |  |
| Fasl |  |

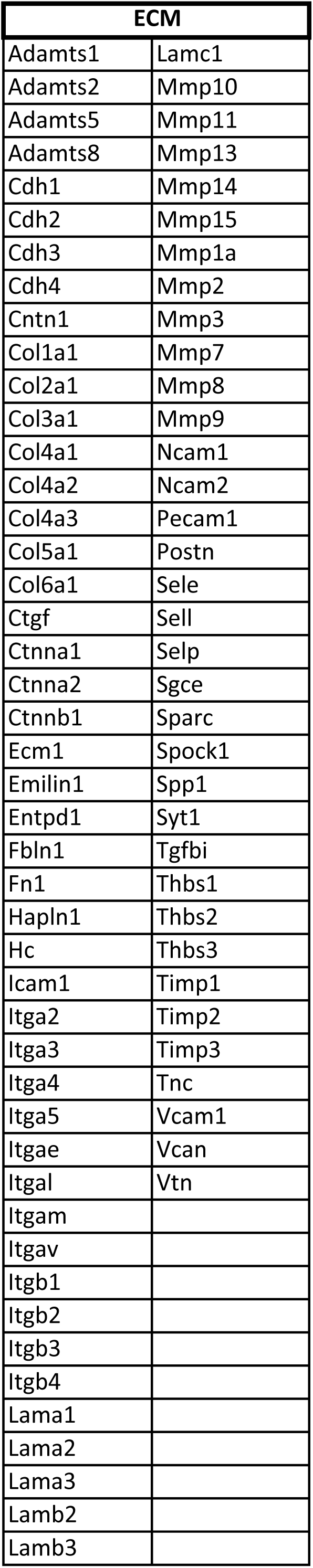

| ROS |  |  |  |
| --- | --- | --- | --- |
| Abl1 | Foxo1 | Ncf1 | Ripk1 |
| Adam9 | Foxo3 | Net1 | Rnf112 |
| Adprhl2 | Foxp1 | Nfe2l2 | Rnf146 |
| Agap3 | Fxn | Ngb | Romo1 |
| Akr1b3 | Fyn | Nme5 | Rps3 |
| Akr1c18 | Gch1 | Nme8 | Sesn1 |
| Akt1 | Gja3 | Nol3 | Sesn2 |
| Aldh3a2 | Glrx | Nos3 | Sesn3 |
| Ankzf1 | Glrx2 | Nox1 | Setx |
| Anxa1 | Gpr37 | Nqo1 | Sigmar1 |
| Apex1 | Gpr37l1 | Nr4a3 | Sirt1 |
| Apoa4 | Gpx1 | Nudt15 | Slc4a1 |
| Apod | Gstp1 | Nup93 | Slc8a1 |
| Aptx | Hdac2 | Oser1 | Sod1 |
| Aqp1 | Hdac6 | Parp1 | Sod2 |
| Atp7a | Hgf | Pawr | Src |
| Axl | Hk3 | Pcgf2 | Stat6 |
| Bcar1 | Hmox1 | Pdcd10 | Stk24 |
| Btk | Hnrnpd | Pde8a | Stk25 |
| Casp3 | Hp | Pdgfra | Stk26 |
| Casp6 | Hsf1 | Pdk2 | Szt2 |
| Cat | Hsph1 | Pink1 | Tacr1 |
| Ccr7 | Il10 | Pld2 | Thbs1 |
| Cd36 | Impact | Plekha1 | Tnfaip3 |
| Cdkn2a | Itgav | Plk3 | Top2b |
| Cfl1 | Jun | Ppargc1b | Traf2 |
| Cflar | Kcnc2 | Ppif | Trap1 |
| Crk | Kdm6b | Ppp1r15b | Trp53 |
| Cryab | Kpna4 | Ppp2cb | Trpa1 |
| Cyp11a1 | Lck | Prdx1 | Trpc6 |
| Cyp1b1 | Lig1 | Prdx2 | Trpm2 |
| Dhfr | Lrrk2 | Prdx3 | Txn1 |
| Ect2 | Map3k5 | Prdx5 | Txndc2 |
| Egfr | Mapk1 | Prdx6 | Txnrd1 |
| Egln1 | Mapk13 | Prkaa1 | Txnrd2 |
| Endog | Mapk3 | Prkca | Txnrd3 |
| Epor | Mapk7 | Prkcd | Ucp1 |
| Ercc6 | Mapk8 | Psap | Ucp2 |
| Ercc6l2 | Mapk9 | Ptk2b | Ucp3 |
| Ezh2 | Mb | Ptprk | Zfp277 |
| Fabp1 | Met | Ptprn | Zfp580 |
| Fancc | Mmp2 | Pycr1 |  |
| Fbln5 | Mmp9 | Reg3b |  |
| Fbxo7 | Mpo | Rela |  |
| Fer | Mpv17 | Rgn |  |
| Fos | Mtr | Rhob |  |

| SASP |  |
| --- | --- |
| Ang5 | Pigf |
| Areg | Plaur |
| Ccl11 | Serpinb2 |
| Ccl12 | Serpine1 |
| Ccl2 | Timp1 |
| Ccl20 | Timp2 |
| Ccl7 | Tnfrsf11b |
| Csf2 | Tnfrsf1a |
| Csf3 | Tnfrsf1b |
| Ctsb | Tnfrsf22 |
| Cxcl11 |  |
| Cxcl12 |  |
| Cxcl13 |  |
| Egf |  |
| Egfr |  |
| Ereg |  |
| Fas |  |
| Fgf2 |  |
| Fgf7 |  |
| Fn1 |  |
| Hgf |  |
| Icam1 |  |
| Ifng |  |
| Igfbp2 |  |
| Igfbp3 |  |
| Igfbp4 |  |
| Igfbp5 |  |
| Igfbp6 |  |
| Igfbp7 |  |
| Il13 |  |
| Il15 |  |
| Il1a |  |
| Il1b |  |
| Il6 |  |
| Il6st |  |
| Il7 |  |
| Mip |  |
| Mmp12 |  |
| Mmp13 |  |
| Mmp14 |  |
| Mmp1a |  |
| Mmp2 |  |
| Mmp3 |  |
| Mmp7 |  |
| Mmp9 |  |
| Ngf |  |

| Senescence |  |
| --- | --- |
| Abl1 | Map2k6 |
| Akt1 | Mdm2 |
| Aldh1a3 | Morc3 |
| Atm | Myc |
| Bmi1 | Nbn |
| Calr | Nfkb1 |
| Ccna2 | Nox4 |
| Ccnb1 | Pcna |
| Ccnd1 | Pik3ca |
| Ccne1 | Plau |
| Cdc25c | Prkcd |
| Cdk2 | Pten |
| Cdk4 | Rb1 |
| Cdk6 | Rbl1 |
| Cdkn1a | Rbl2 |
| Cdkn1b | Serpinb2 |
| Cdkn1c | Serpine1 |
| Cdkn2a | Sirt1 |
| Cdkn2b | Sod1 |
| Cdkn2c | Sod2 |
| Cdkn2d | Sparc |
| Chek1 | Tbx2 |
| Chek2 | Tbx3 |
| Cited2 | Terf2 |
| Col1a1 | Tert |
| Col3a1 | Tgfb1 |
| Creg1 | Tgfb1i1 |
| E2f1 | Thbs1 |
| E2f3 | Trp53 |
| Egr1 | Trp53bp1 |
| Ets1 | Twist1 |
| Ets2 | Vim |
| **Fn1** |  |
| Gadd45a |  |
| Glb1 |  |
| Gsk3b |  |
| Hras |  |
| Id1 |  |
| Igf1 |  |
| Igf1r |  |
| Igfbp3 |  |
| Igfbp5 |  |
| Igfbp7 |  |
| Ing1 |  |
| Map2k1 |  |
| Map2k3 |  |

